# The metabolic repair enzyme phosphoglycolate phosphatase regulates central carbon metabolism and fosmidomycin sensitivity in *Plasmodium falciparum*

**DOI:** 10.1101/415505

**Authors:** Laure Dumont, Mark B Richardson, Phillip van der Peet, Matthew WA Dixon, Spencer J Williams, Malcolm J McConville, Leann Tilley, Simon A Cobbold

## Abstract

The asexual blood stages of the malaria parasite, *Plasmodium falciparum* are highly dependent on glycolysis for ATP synthesis, redox balance and provision of essential anabolic precursors. Recent studies have suggested that members of the haloacid dehalogenase (HAD) family of metabolite phosphatases may play an important role in regulating multiple pathways in *P. falciparum* central carbon metabolism. Here, we show that the *P. falciparum* HAD protein, phosphoglycolate phosphatase (*Pf*PGP), which is homologous to yeast Pho13 and mammalian PGP, regulates glycolysis in asexual blood stages by controlling intracellular levels of several intermediates and novel end-products of this pathway. Deletion of the *P. falciparum pgp* gene significantly attenuated asexual parasite growth in red blood cells, while comprehensive metabolomic analysis revealed the accumulation of two previously uncharacterized metabolites, as well as changes in a number of intermediates in glycolysis and the pentose phosphate pathway. The two unknown metabolites were assigned as 2-phospho-lactate and 4-phosphoerythronate by comparison of their mass spectra with synthetic standards. 2-Phospho-lactate was significantly elevated in wildtype and Δ*Pf*PGP parasites cultivated in the presence of methylglyoxal and D-lactate, but not L-lactate, indicating that it is a novel end-product of the methylglyoxal pathway. 4-Phosphoerythronate is a putative side product of the glycolytic enzyme, glyceraldehyde dehydrogenase and the accumulation of both 4-phosphoerythronate and 2-phospho-D-lactate were associated with changes in glycolytic and the pentose phosphate pathway fluxes as shown by ^13^C-glucose labelling studies and increased sensitivity of the Δ*Pf*PGP parasites to the drug fosmidomycin. Our results suggest that *Pf*PGP contributes to a novel futile metabolic cycle involving the phosphorylation/dephosphorylation of D-lactate as well as detoxification of metabolites, such as 4-phosphoerythronate, and both may have important roles in regulating *P. falciparum* central carbon metabolism.

**Author summary:** The major pathogenic stages of the malaria parasite, *Plasmodium falciparum*, develop in red blood cells where they have access to an abundant supply of glucose. Unsurprisingly these parasite stages are addicted to using glucose, which is catabolized in the glycolytic and the pentose phosphate pathways. While these pathways also exist in host cells, there is increasing evidence that *P. falciparum* has evolved novel ways for regulating glucose metabolism that could be targeted by next-generation of anti-malarial drugs. In this study, we show the red blood cell stages of *P. falciparum* express an enzyme that is specifically involved in regulating the intracellular levels of two metabolites that are novel end-products or side products of glycolysis. Parasite mutants lacking this enzyme are viable but exhibit diminished growth rates in red blood cells. These mutant lines accumulate the two metabolites, and exhibit global changes in central carbon metabolism. Our findings suggest that metabolic end/side products of glycolysis directly regulate the metabolism of these parasites, and that the intracellular levels of these are tightly controlled by previously uncharacterized metabolite phosphatases.

## Introduction

*Plasmodium falciparum* is the major cause of malaria, a disease that continues to kill ∼ 445,000 people each year and has significant impacts on the development of some of the poorest countries (1). The symptoms of malaria arise from the progressive ∼ 48-hour cycles of parasite invasion into host red blood cells (RBC), rapid growth of asexual parasite stages within RBC and subsequent RBC lysis. RBC provide intracellular parasite stages with abundant supplies of glucose, amino acids and other carbon sources derived from the serum and/or breakdown of RBC proteins and lipids. *P. falciparum* asexual blood stages primarily use glucose as their major carbon source and are largely dependent on glycolysis for generation of ATP, although they retain a low flux TCA cycle for generation of the mitochondrial membrane potential (2-4). Rates of glucose utilization and L-lactate production from glycolysis are increased up to 100-fold in *P. falciparum*-infected RBC compared to uninfected RBC (5). The high glycolytic flux of *P. falciparum* asexual stages provides these cells with sufficient ATP and biosynthetic precursors to sustain anabolic processes and high rates of replication. However, high rates of glycolysis can generate toxic metabolic end-products, such as methylglyoxal, which can lead to increased oxidative stress, chemical modification and denaturation/inactivation of proteins, lipids and DNA (6-8). Thus, it is likely that *P. falciparum* has evolved ways to regulate pathways such as glycolysis under nutrient excess conditions.

*P. falciparum* parasites express only a limited number of transcription factors and possess limited nutrient-stimulated transcriptional control (9-12), indicating that regulation of central carbon metabolism primarily occurs at the post-transcriptional level. It is intriguing that biochemical investigations have uncovered a lack of conventional eukaryote allosteric regulatory/feedback mechanisms, suggesting dependence on other post-transcriptional regulatory mechanisms (13, 14). One group of proteins that has a role in regulating glycolysis in *P. falciparum* asexual stages is the haloacid dehalogenase (HAD) family of metabolite phosphatases. The first member of this family to be functionally characterized was *Pf*HAD1, identified in a screen for *P. falciparum* mutant lines that were resistant to the isoprenoid biosynthesis inhibitor fosmidomycin (15). HAD1 exhibited broad *in vitro* phosphatase activity against a range of sugar-phosphates and triose-phosphates that are connected to glycolysis, suggesting that it might have a role *in vivo* in the promiscuous dephosphorylation of glycolytic intermediates and negatively regulate glycolytic fluxes. Mutational inactivation of HAD1 leads to increased glycolytic flux and flow of intermediates into anabolic pathways such as isoprenoid biosynthesis, with associated increase in fosmidomycin resistance. Interestingly, a second HAD enzyme, HAD2 was identified in the same screen and was shown to exhibit *in vitro* phosphatase activity against glycolytic intermediates, suggesting that HAD enzymes may regulate multiple pathways in *P. falciparum* central carbon metabolism (16).

One of the HAD family members in the *P. falciparum* genome is homologous to the enzyme phosphoglycolate phosphatase (PGP), which is involved in regulating intracellular levels of several metabolites, including glycerol-3-phosphate (Gro3P), 2-phosphoglycolate, 2-phospho-L-lactate and 4-phosphoerythronate (4-PE) (17, 18). 2-Phospho-L-lactate and 4-PE are thought to be minor side-products of the high flux reactions catalyzed by pyruvate kinase and glyceraldehyde-3-phosphate dehydrogenase (GAPDH), respectively (17). PGP may thus act as a metabolite repair enzyme that detoxifies metabolites that would otherwise accumulate and allosterically affect key enzymes in central carbon metabolism (19, 20). In this study we have investigated the role of *P. falciparum* PGP in asexual blood stages. In contrast to the situation in animal cells, we find that *P. falciparum* accumulates 2-phospho-D-lactate, rather than 2-phospho-L-lactate, indicating synthesis through the methylglyoxal pathway rather than via enzymes in lower glycolysis. We show that *Pf*PGP regulates the level of this novel end-product and contributes to a metabolic futile cycle that may regulate intracellular ATP levels. *Pf*PGP is also involved in detoxifying 4-PE and we provide evidence that the accumulation of 4-PE leads to dysregulation of the pentose phosphate pathway and glycolysis, as well as increased sensitivity to fosmidomycin. Overall, these data highlight novel aspects of *P. falciparum* glycolysis and a key role for *Pf*PGP in regulating central carbon metabolism in asexual blood stages.

## Results

### *P. falciparum* PGP is a cytoplasmic protein required for normal growth of asexual stages in RBC

The *P. falciparum* gene, PF3D7_0715000 (hereinafter referred to as *Pf*PGP) shares homology to the phosphoglycolate phosphatase (PGP) family of HAD enzymes that are involved in regulating the intracellular levels of key phospho-intermediates or end-products generated in glycolytically active cells, and features all four characteristic HAD motifs in its sequence (Fig. S1) (21). Previous studies have suggested that *Pf*PGP may be involved in vitamin B1 biosynthesis, although this role has yet to be confirmed (22). Given the strong dependence of *P. falciparum* asexual blood stages on glycolysis for generation of ATP, redox balance and generation of anabolic precursors, we reinvestigated the functional role of *Pf*PGP. Consistent with previous studies (22), we show that a *Pf*PGP-GFP fusion protein is exclusively located in the cytoplasm of asexual blood stages (Fig. 1a) and is readily extracted in either phosphate buffer saline/sodium dodecyl sulfate (SDS) or radioimmunoprecipitation assay (RIPA) buffer/SDS (Fig. S2a).

**Figure 1.**
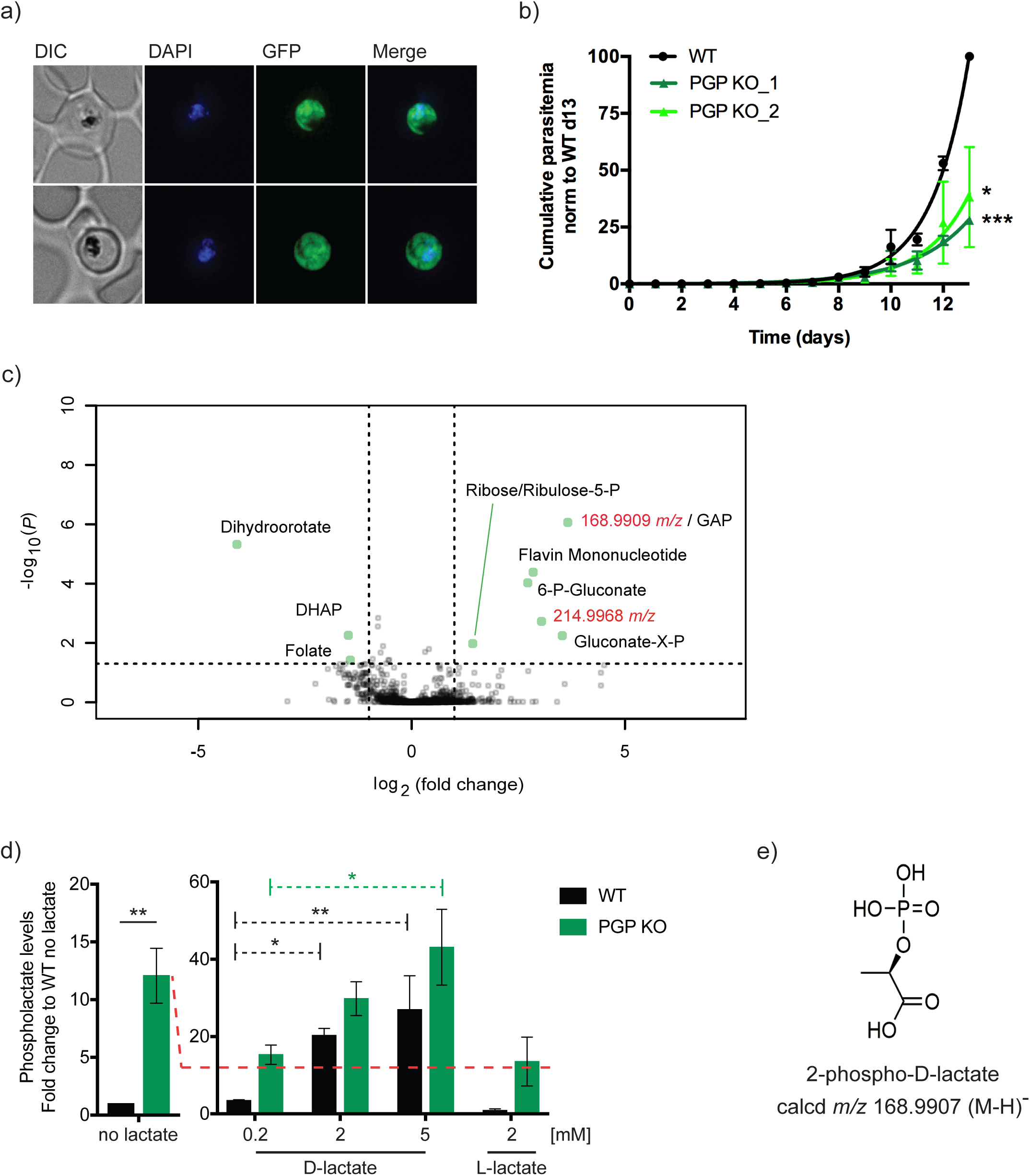
*Pf*PGP is required for normal growth of *P. falciparum* asexual stages and Δ*Pf*PGP mutant parasites selectively accumulate metabolites in the PPP, as well as two novel metabolites including 2-phospholactate. a) Fluorescence microscopy of live *Pf*PGP-GFP-infected RBC. Infected RBC were labelled with DAPI and visualized by differential interference contrast (DIC) and fluorescence microscopy. GFP fluorescence was localized throughout the cytoplasm. b) The asexual growth of Δ*Pf*PGP versus wildtype (WT) infected RBC was monitored daily over a 13-day period by flow cytometry following SYTO61 labelling. Data are presented as the mean ± SEM of the cumulative parasitemia normalised to the WT day 13 data point (100 %), from three independent experiments. Statistical significance was determined using a paired t-test at day 13 (* and *** denote *P* values < 0.05 and < 0.001 respectively). c) WT and Δ*Pf*PGP-infected RBC were harvested, and intracellular metabolite levels profiled by GC-MS. A volcano plot of change log_2_ (fold-change) versus significance –log_10_ (*P*) for the Δ*Pf*PGP mutant parasites compared to wildtype parasites. Annotated metabolites were verified using standards with the exception of gluconate-X-P (where the position of the P is unknown). Ribose-5-P and ribulose-5-P co-elute under the chromatography conditions used. Unknown metabolites with 214.9968 *m/z*, and 168.9909 *m/z* increased in the mutant line. d) WT- and Δ*Pf*PGP-infected RBC were incubated with different concentrations of L- or D-lactate and intracellular levels of phospholactate measured by GC-MS. Data are presented as fold-change versus the WT - no lactate condition. Data are presented as the mean ± SEM from three independent experiments performed on different days. Statistical significance was determined using unpaired t-testing for the no lactate condition WT vs Δ*Pf*PGP (** denotes a *P* value <0.01); and one-way ANOVA for all lactate-stimulation conditions (dotted lines; WT: * and ** denote *P* values < 0.05 and 0.01, respectively; PGP KO: * denotes a *P* value <0.05; all other comparisons were non-significant). e) Structure of 2-phospho-D-lactate (C_3_H_7_O_6_P).

To characterise the role of *Pf*PGP *in vivo*, a knock-out parasite line was generated using CRISPR/Cas9 (Fig. S2b) (23). PGP-deleted *P. falciparum* mutants (Δ*Pf*PGP) were recovered and loss of the gene was confirmed by PCR (Fig. S2c). Analysis of the growth of two clones of the Δ*Pf*PGP line indicated that loss of this gene was associated with a significant decrease in growth rate (Fig. 1b; *P* = 0.0005 and 0.039 for clones 1 and 2, respectively). While the NF54 parental line had a doubling time of 1.04 ± 0.12 days, Δ*Pf*PGP clones 1 and 2 had doubling times of 1.48 ± 0.24 days and 1.21 ± 0.03 days, respectively. As the culture medium contains vitamin B1 and other vitamins, these data support the premise that *Pf*PGP has a metabolic function distinct from vitamin B1 biosynthesis.

To define the function of *Pf*PGP, we undertook a comprehensive metabolomic analysis of parental (NF54) and Δ*Pf*PGP mutant parasite lines. Trophozoite-stage infected RBC were magnetically enriched (> 95% parasitemia), and metabolites were extracted and analyzed by liquid chromatography - mass spectrometry (LC-MS). The mutant parasite line showed selective increases in a number of metabolites associated with the pentose phosphate pathway (6-phosphogluconate, ribose/ribulose-5-phosphate), flavin mononucleotide and two unknown peaks with accurate masses (*m/z*; mass over charge ratio) of 168.9909 and 214.9968, respectively (Fig. 1c). Candidate metabolites for 168.9909 *m/z* included the glycolytic intermediates, glyceraldehyde phosphate (GAP) and dihydroxyacetone phosphate (DHAP), or the two enantiomers of phospho-D/L-lactate (METLIN metabolite database). GAP or DHAP were discounted based on the lack of co-elution with authentic standards for these metabolites on LC-MS or gas chromatography – mass spectrometry (GC-MS), indicating that the 168.9909 *m/z* peak may be phospho-D/L-lactate. To distinguish between the two possible enantiomers, parasite-infected RBC were incubated in the presence of either L- or D-lactate and intracellular levels of the 168.9909 *m/z* peak determined by targeted GC-MS analysis (Fig. 1d). Consistent with the untargeted analysis, accumulation of 168.9909 *m/z* was increased 12-fold (± 2.4) in the Δ*Pf*PGP line compared to the parental wildtype (WT) line (Fig. 1d, left panel). Addition of 2 mM L-lactate (the major enantiomer generated by glycolysis) to WT or Δ*Pf*PGP parasite cultures had no effect on the intracellular levels of this metabolite. In contrast, incubation of either WT or Δ*Pf*PGP parasites with D-lactate resulted in a marked increase of the 168.9909 *m/z* peak (Fig. 1d, right panel). The 168.9909 *m/z* peak was confirmed as phospholactate by synthesizing racemic 2-phospho-D/L-lactate. This standard had the same fragmentation profile and retention time as the peak of interest via GC-MS (Fig. S3). Taken together, these data indicate that WT parasites are capable of phosphorylating endogenous and exogenous D-lactate to form 2-phospho-D-lactate (Fig. 1e), and that *Pf*PGP is involved in dephosphorylating this species back to D-lactate.

### Phospholactate is a product of the methylglyoxal pathway

In animal cells, 2-phospho-L-lactate is thought to be a by-product of the terminal glycolytic enzyme, pyruvate kinase (17). The finding that *P. falciparum* phospholactate is derived from the D-stereoisomer of lactate indicates, for this case, that an additional pathway for synthesizing this intermediate must operate. The only pathway known to generate D-lactate in *P. falciparum* is the methylglyoxal pathway, which is required for detoxification of methylglyoxal formed by non-enzymatic phosphate elimination of the glycolytic triose phosphates, GAP and DHAP (7, 24). Methylglyoxal is converted to *S*-D-lactoyl-glutathione by glyoxalase I (GloI) and then further metabolized to D-lactate by glyoxalase II (GloII) (6) (Fig. 2a). To investigate whether the D-lactate generated in this pathway is converted to 2-phospho-D-lactate we generated *P. falciparum* knock-out lines lacking the cytoplasmic *gloI* gene (PF3D7_1113700) using the CRISPR/Cas9 system (Fig. S4a) (23). Knock-out of *PfgloI* was confirmed by PCR (Fig. S4b). Analysis of lysates of saponin-purified *P. falciparum* trophozoites indicated that the ΔGloI line had greatly reduced GloI activity *in vitro*, as measured by conversion of methylglyoxal and glutathione to D-lactate by GC-MS (Fig. 2b, Table S1). Specifically, the ΔGloI parasites exhibited an 80% reduction in D-lactate production after 60 minutes (WT = 100 %, GloI KO = 20.61 % ± 9.1). Loss of *Pf*GloI was associated with a statistically significant growth defect over 13 days (Fig. 2c; GloI KO_1 *P* = 0.0025 and GloI KO_2 *P* = 0.038; doubling time WT = 1.04 ± 0.12 days / GloI KO_1 = 1.40 ± 0.40 days / GloI KO_2 = 1.26 ± 0.27 days) indicating that detoxification of methylglyoxal is important for normal asexual growth and development in RBC.

**Figure 2.**
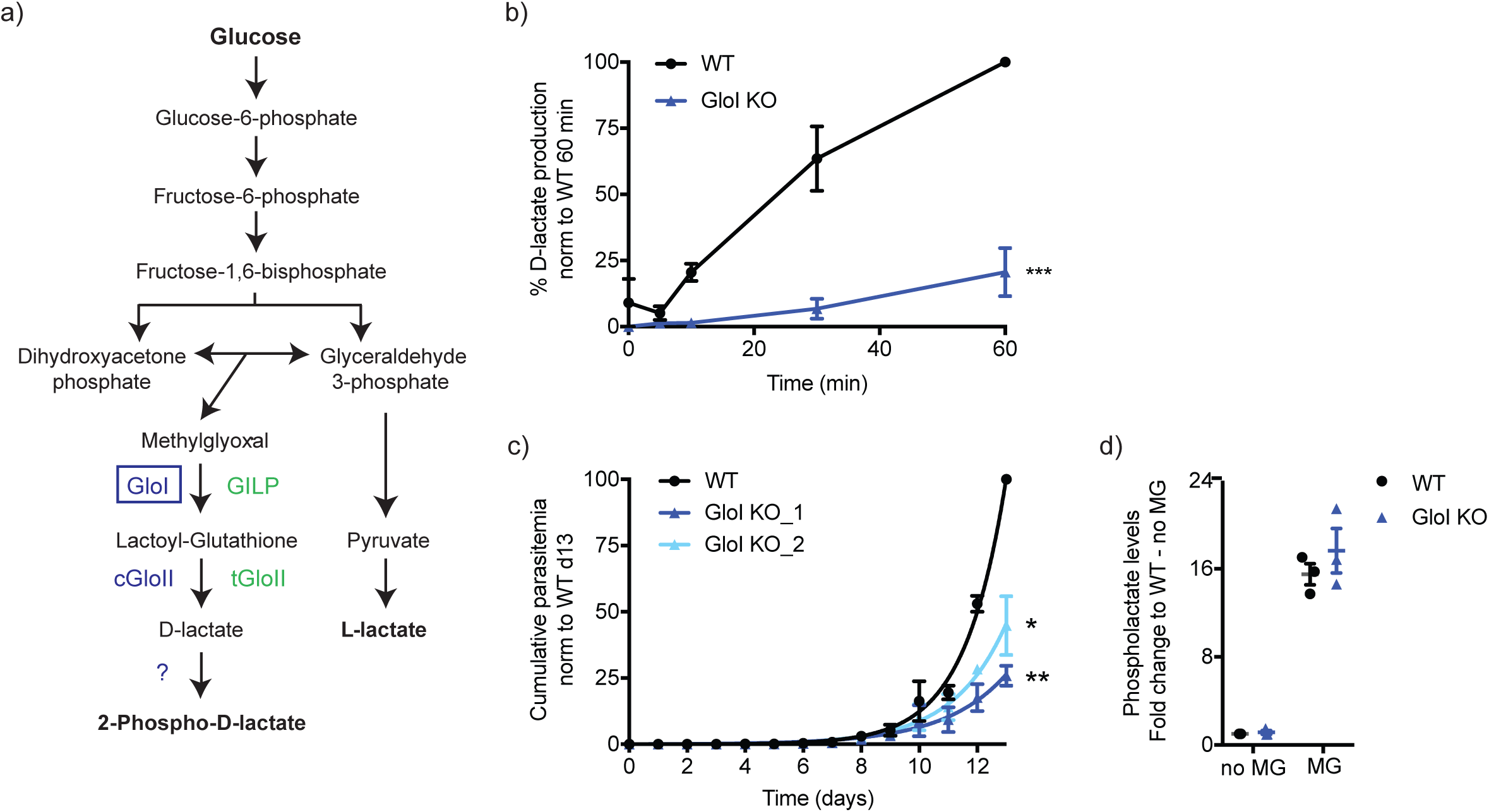
2-Phospho-D-lactate is generated via the glyoxalase pathway. a) Schematic of the methylglyoxal detoxification system in *P. falciparum*. GloI and cGloII catalyze the conversion of the toxic glycolytic overflow metabolite, methylglyoxal, to D-lactate. *P. falciparum* also expresses GloI-like protein (GILP) and tGloII which are thought to be targeted to the apicoplast. b) *in vitro* production of D-lactate in ΔGloI parasites versus WT parasites. Data are presented as the % of D-lactate production normalised to the WT 60-minute time point (100 %) after experimental background subtraction. Data are presented as the mean ± SEM from three independent experiments performed on different days and statistical significance was determined using unpaired t-testing at the 60-minute time point (*** denotes a *P* value < 0.001). c) The asexual growth of ΔGloI-versus WT-infected RBC was monitored daily over a 13-day period by flow cytometry following SYTO 61 labelling. Results are presented as the cumulative parasitemia normalised to the WT day 13 data point (100%), taking into account the dilutions made each cycle. Data are presented as the mean ± SEM from three independent experiments performed on different days and statistical significance was determined using paired t-testing at day 13 (* and ** denote *P* values <0.05 and 0.01 respectively). d) WT- and ΔGloI-infected RBC were incubated with no or 1mM methylglyoxal (MG) and intracellular levels of 2-phospholactate levels were measured by GC-MS. Data are presented as fold change to the WT - no methylglyoxal condition. Data are presented as the mean ± SEM from three independent experiments performed on different days.

To investigate whether 2-phospho-D-lactate is synthesized by the methylglyoxal pathway, parental WT and ΔGloI parasite cultures were suspended in medium containing methylglyoxal (1 mM, 1 hour, 37°C), then levels of 2-phospho-D-lactate were measured by GC-MS. Addition of methylglyoxal led to a 15-fold increase in 2-phospho-D-lactate levels, indicating that this metabolite is the end-product of the methylglyoxal pathway (Fig. 2d). Strikingly, similar levels of 2-phospho-D-lactate were present in both WT and ΔGloI parasites, before and after addition of methylglyoxal, indicating that synthesis of D-lactate is not strictly dependent on *Pf*GloI. It is possible that the apicoplast isoform of GloI (GloI-like protein, GILP) could substitute for cytoplasmic GloI and/or that methylglyoxal can be converted to *S*-D-lactoyl-glutathione non-enzymatically. Alternatively, methylglyoxal may be converted to D-lactate by the RBC methylglyoxal pathway and D-lactate subsequently imported by the parasite and converted to 2-phospho-D-lactate. Collectively, these studies suggest that 2-phospho-D-lactate is generated by the direct phosphorylation of D-lactate produced by either the host cell, or the parasite methylglyoxal detoxification pathways.

### Accumulation of 2-phospho-D-lactate is toxic at non-physiological concentrations

Next, we determined whether the reduced growth rate of the Δ*Pf*PGP mutant lines in RBC could be attributed to the toxic accumulation of 2-phospho-D-lactate. WT- and Δ*Pf*PGP-infected RBC were suspended in medium containing 0, 1 and 5 mM D-lactate to elevate intracellular levels of 2-phospho-D-lactate and asexual parasite growth was monitored over a period of 13 days. Growth of both WT and Δ*Pf*PGP, were significantly reduced in the presence of a high (5 mM) concentration of D-lactate (Fig. 3a,b), with Δ*Pf*PGP appearing to be more susceptible to the metabolic treatment (growth decreased by 49.49% ± 2.31 compared to 20.25% ± 5.24 for WT). This result suggests that accumulation of 2-phospho-D-lactate is toxic to the parasite and that *Pf*PGP plays a key role in maintaining non-toxic levels. However, extracellular concentrations of D-lactate in *P. falciparum* cultures are generally below 1 mM indicating that 2-phospho-D-lactate toxicity may not be the major cause of the growth defect of the Δ*Pf*PGP mutant under physiological growth conditions.

**Figure 3.**
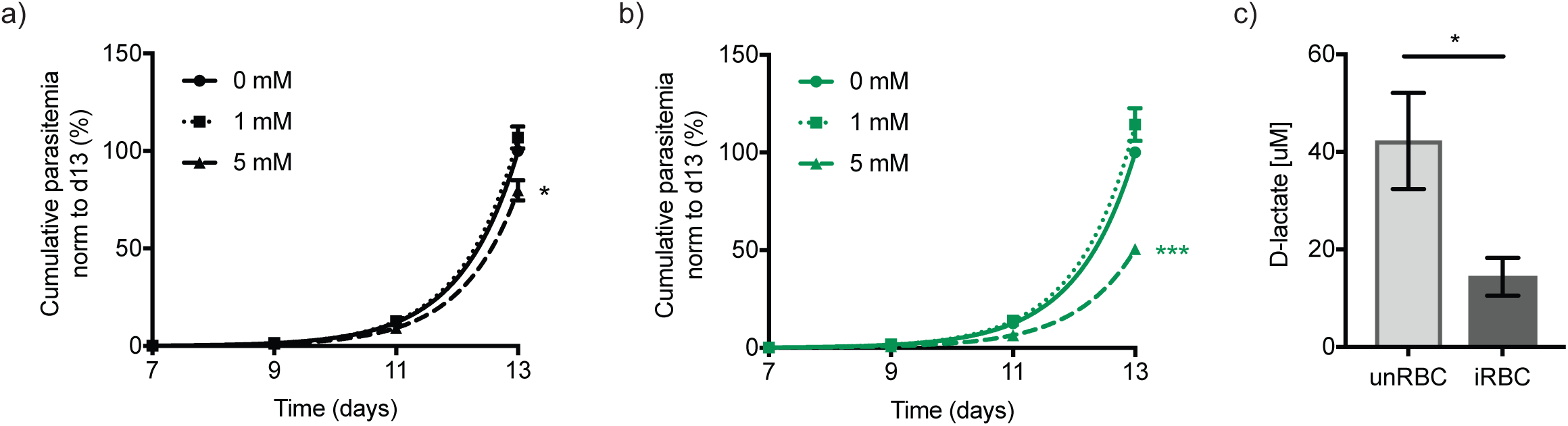
Accumulation of 2-phospho-D-lactate is only toxic at high concentrations. a) WT- and b) Δ*Pf*PGP-infected RBC were incubated with 0, 1 or 5 mM D-lactate and asexual growth was monitored every 48 hours over a 13-day period by flow cytometry following SYTO 61 labelling. Data from four independent experiments performed on different days are presented as the mean ± SEM of the cumulative parasitemia normalised to the 0 mM/ day 13 data point (100%). Dilutions of the cultures were taken into account and statistical significance was determined using paired t-testing at day 13 (* and *** denote *P* < 0.05 and 0.001 respectively). c) Extracellular levels of D-lactate were measured in uninfected RBC (unRBC) and WT-infected RBC (iRBC) after sample deproteinisation using a D-lactate plate assay (Cayman chemicals). Cultures were set at 2.5% haematocrit (and 1.8 to 4% parasitemia for infected RBC). Media was collected after 26-29 hours of culture (ring to trophozoite stages for infected RBC). Data are presented as the mean ± SEM from four independent repeats collected on different days and statistical significance was determined using unpaired t-testing (* denotes *P* < 0.05).

An alternative possibility is that the inter-conversion of D-lactate and 2-phospho-D-lactate, mediated by an unknown kinase and *Pf*PGP, has a metabolic function that is required for parasite asexual growth. We hypothesized that D-lactate may be converted to 2-phospho-D-lactate to prevent its secretion with L-lactate. Consistent with this proposal, analysis of the culture medium of uninfected and infected RBC showed that secretion of D-lactate was reduced by ∼75% in the latter, indicating that D-lactate may be sequestered within the parasite as 2-phospho-D-lactate (Fig. 3c). This finding differed from the previously reported ∼30-fold increase in D-lactate secretion following erythrocyte infection (24). Here we deproteinised samples before detection and suspect that without this step, misleadingly high levels of D-lactate are observed due to residual enzyme activity in the media.

### Loss of *Pf*PGP leads to increased flux through the pentose phosphate pathway

Our untargeted LC-MS metabolomic studies identified a second unknown metabolite accumulating in the Δ*Pf*PGP mutant of 214.9968 *m/z* (Fig. 1c). METLIN database searching suggested that this metabolite might be 4-phosphoerythronate (4-PE; Fig. 4a). 4-PE is not an intermediate in canonical metabolic pathways but is thought to be generated by GAPDH acting on the pentose phosphate pathway intermediate erythrose-4-phosphate, instead of its preferred substrate, GAP (17). The identity of the 214.9968 *m/z* peak was confirmed as 4-PE through comparison of its MS spectrum and GC-MS retention time with those of a synthetically-acquired standard (Fig. 4b and Table S1). The intracellular levels of 4-PE were not affected by addition of exogenous D- or L-lactate to WT- and Δ*Pf*PGP-infected RBC cultures, indicating that this metabolite is generated independently of the D-lactate/ 2-phospho-D-lactate pathway (Fig. S5).

**Figure 4.**
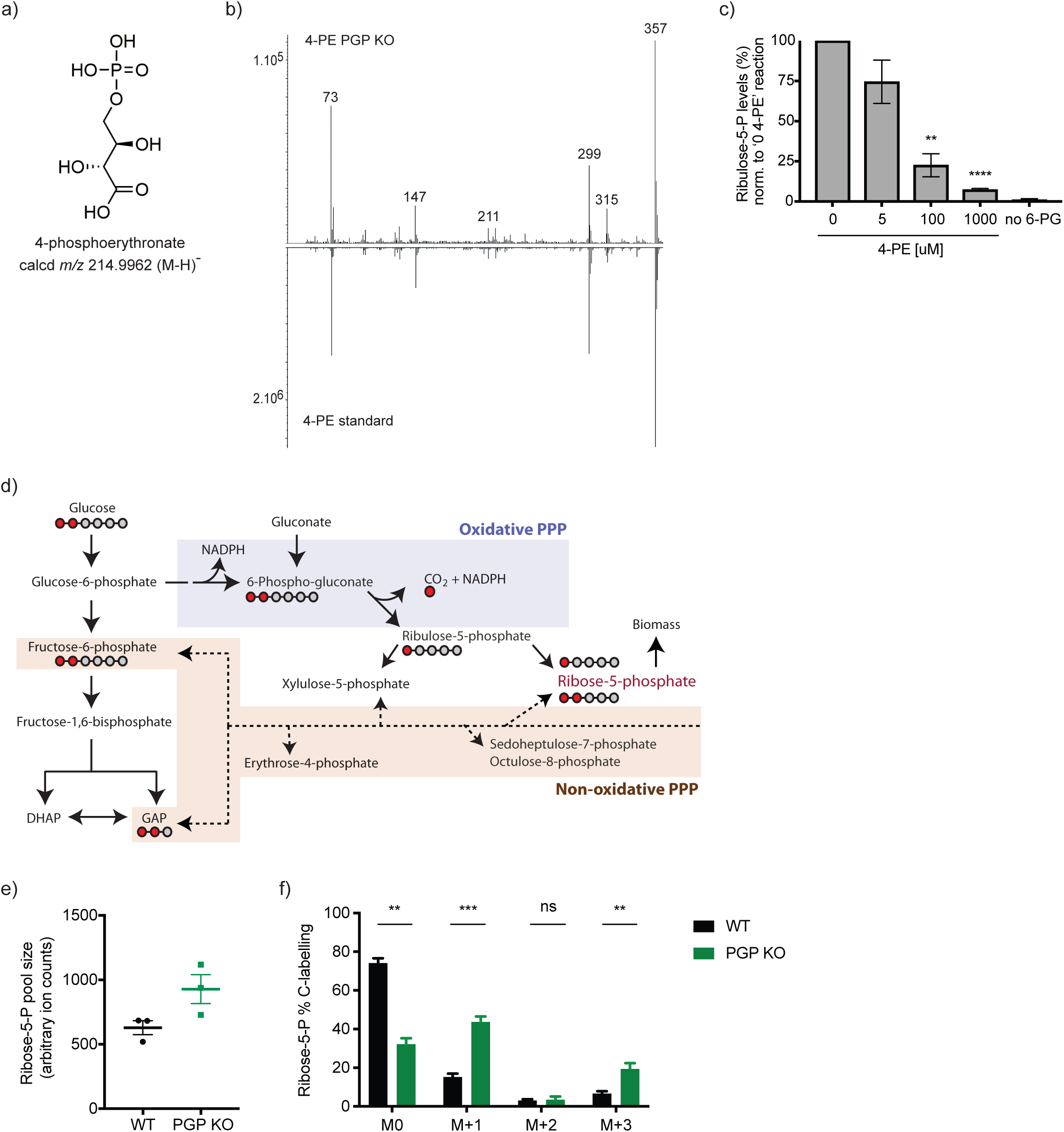
Loss of *Pf*PGP leads to partial inhibition of 6-PGD, and enhanced flux through the oxidative PPP. a) Chemical structure of 4-PE. b) GC-MS mass spectrum of *P. falciparum* 4-PE is identical to chemically synthesised 4-PE. c) Synthetic 4-PE was added to lysates of saponin-purified trophozoites, together with 6-phosphogluconate (6-PG) and activity of 6-PGD assessed by measurement of ribulose-5-P by GC-MS. Data are presented as fold change to the 0, 4-PE condition (100%). Data are presented as the mean ± SEM from four to five independent experiments performed on different days and statistical significance was determined using one-way ANOVA in comparison to the 0, 4-PE condition (** and **** denote *P* < 0.01 and *P* < 0.0001 respectively). d) Schematic of the pentose phosphate pathway in *P. falciparum*. Where relevant, carbon backbones are presented as grey (unlabelled) or red (^13^C-labelled) dots. d-e) The relative activities of the oxidative and non-oxidative pentose phosphate pathways was monitored with ^13^C-1,2-glucose incorporation into ribose-5-P. The ribose-5-P pool sizes were evaluated (e) as well as the labelling pattern (f). M+1 represents the fraction of ribose-5-P derived from the oxidative arm, M+2 represents the contribution of the non-oxidative arm, and M+3 comprises both oxidative and non-oxidative labelling. Data are presented as the mean ± SEM from three independent experiments performed on different days. Statistical significance was determined using unpaired t-testing (* and *** denote *P* < 0.01 and *P* <0.001 respectively).

4-PE is an inhibitor or allosteric regulator of enzymes in the pentose phosphate pathway (PPP), including 6-phospho-gluconate dehydrogenase (6-PGD) (17, 25). Consistent with this fact, we observed an increase in 6-phosphogluconate, the substrate for 6-PGD, in the Δ*Pf*PGP parasites (Fig. 1c). To confirm that 4-PE inhibits 6-PGD directly, we measured the *in vitro* activity of 6-PGD in lysates of saponin-lysed trophozoite-infected RBC in the absence and presence of 4-PE. Lysates were incubated with 6-phosphogluconate and increasing concentrations of 4-PE and the conversion of 6-phosphogluconate to ribulose-5-P was assayed by GC-MS (Table S1). Partial inhibition was observed between 5 - 100 μM and complete inhibition at 1 mM 4-PE (Fig. 4c). Neither erythronate nor 2-phospholactate exhibited inhibitory effects in this assay (Fig. S6). These results suggest that the accumulation of 4-PE in Δ*Pf*PGP parasites (μM range) may partially inhibit activity of 6-PGD *in vivo*.

Paradoxically, our metabolomic studies indicated that levels of downstream intermediates in the oxidative and non-oxidative PPP (ribose-5-P/ribulose-5-P) were elevated, rather than decreased, in Δ*Pf*PGP parasites (Fig. 1c). An increase in the pool size of these pentose-phosphates could reflect increased flux through the non-oxidative PPP and/or a compensating increase in flux through the oxidative PPP (overcoming the partial inhibition of 6-PGD). To address this question directly, enriched WT- and Δ*Pf*PGP-infected RBC cultures were metabolically labelled with ^13^C-1,2-glucose to measure fluxes through both arms of the PPP. Catabolism of ^13^C-1,2-glucose through the oxidative arm of the PPP leads to loss of ^13^C on carbon-1 of 6-phosphogluconate (as carbon dioxide) as this intermediate is converted to ^13^C_1_-ribose-5-P. In contrast, conversion of ^13^C-1,2-glucose to ribose-5-P via the non-oxidative pathway does not involve a decarboxylation step resulting in ^13^C_2_-ribose-5-P (Fig. 4d). Purified WT- and Δ*Pf*PGP-infected RBC were incubated with ^13^C-1,2-glucose for 30 minutes at 37°C and ^13^C-enrichment in ribose-5-P was determined by GC-MS. Ribose-5-P levels were elevated in Δ*Pf*PGP parasite cultures (Fig. 4e, Table S1), consistent with the results of the initial metabolomic analyses (Fig. 1c). The overall rate of turnover of pentose-phosphates were significantly increased in the Δ*Pf*PGP mutant (Fig. 4f, M0 fraction). This was entirely due to increased production of ^13^C_1_-ribose-5-P (and ^13^C_3_-ribose-5-P) and indicates an increased flux through the oxidative PPP. It is notable that the flux through the non-oxidative PPP (as indicated by levels of ^13^C_2_-ribose-5-P) was relatively low in both WT and Δ*Pf*PGP parasites (Fig. 4f). These results suggest that inhibition of 6-PGD in the Δ*Pf*PGP parasite lines, due to the accumulation of 4-PE, is more than compensated for by increased flux through this pathway. Therefore, partial inhibition of the oxidative PPP in the Δ*Pf*PGP line is an unlikely cause of the decreased growth rate of asexual stages in RBC.

### Loss of *Pf*PGP is associated with changes in glycolytic flux and increased sensitivity to fosmidomycin

The untargeted LC-MS profiling of Δ*Pf*PGP parasites revealed that DHAP was one of the few metabolites to be significantly down-regulated in the mutant (Fig. 1c). The interconversion of DHAP and GAP is mediated by the glycolytic enzyme, triosephosphate isomerase (TPI) which, like 6-PGD, catalyzes a reaction utilizing an ene-diolate intermediate that might be sensitive to allosteric modulation by 4-PE (26, 27). DHAP and other triose-phosphates are catabolyzed in the glycolytic pathway and imported into the apicoplast where they are used for synthesis of 1-deoxy-D-xylulose-5-P, the first committed intermediate in isoprenoid biosynthesis. Conversion of 1-deoxy-D-xylulose-5-P into 2-*C*-methyl-D-erythritol-4-P is inhibited by fosmidomycin, a potent antimalarial (Fig. 5a) (28). We hypothesised that perturbation of glycolytic flux and/or balance of triose-phosphates in the Δ*Pf*PGP mutant could lead to reduced isoprenoid biosynthesis and increased sensitivity to fosmidomycin. We evaluated the sensitivity of Δ*Pf*PGP-infected RBC to fosmidomycin in a 72-hour drug treatment assay and measured effects upon parasite growth via flow cytometry (Fig. 5b). In comparison to WT, the Δ*Pf*PGP mutant exhibited a significant 4-fold increase in fosmidomycin sensitivity (EC_50_ WT = 358 nM ± 14; Δ*Pf*PGP = 89 nM ± 9.8; *P* = 0.001). These findings suggest that elevated 4-PE levels in Δ*Pf*PGP parasites lead to perturbations in glycolytic flux, reduced isoprenoid synthesis and concomitant increase in fosmidomycin sensitivity.

**Figure 5.**
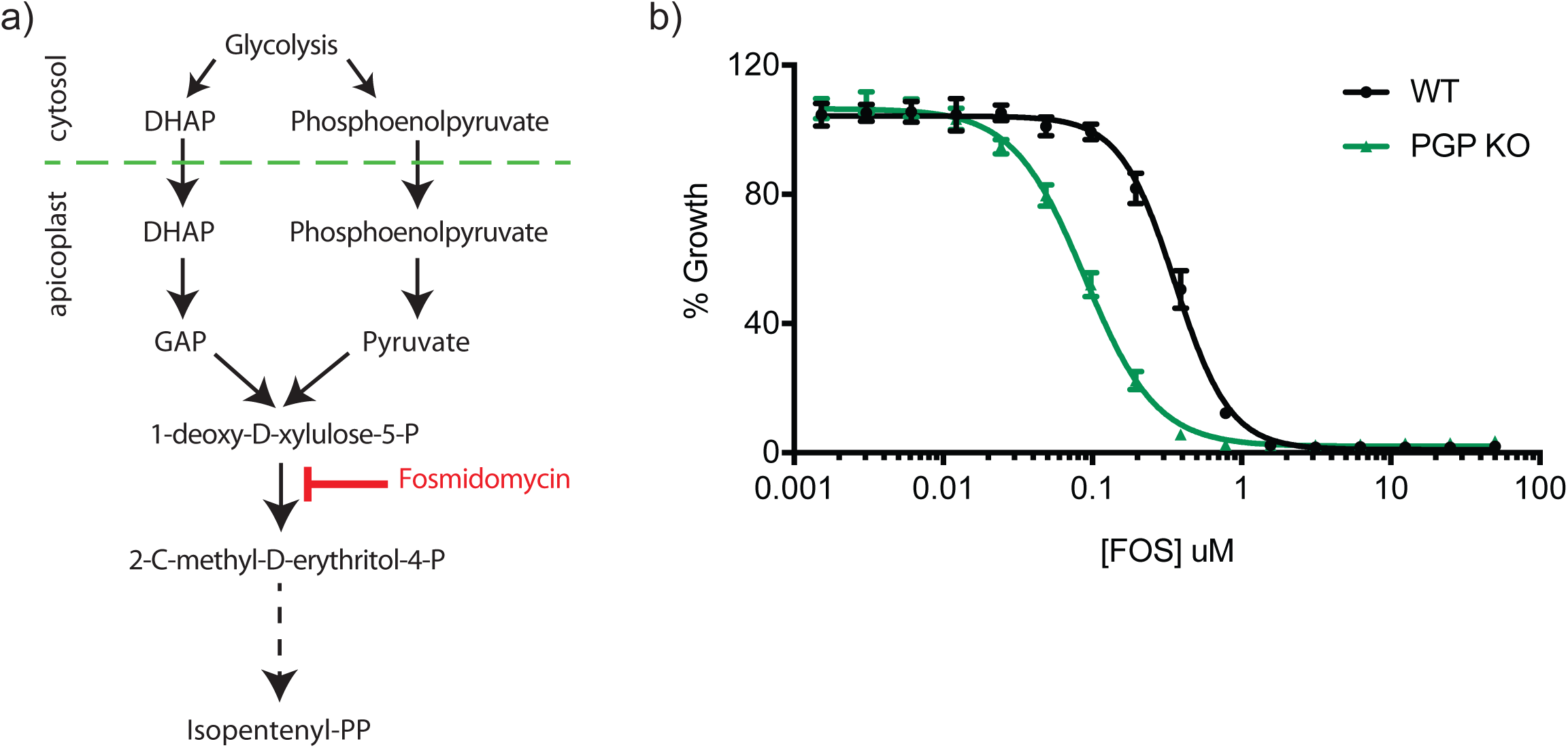
Loss of *Pf*PGP leads to increased sensitivity to fosmidomycin. a) DHAP and phosphoenolpyruvate, derived from glycolysis, are imported into the apicoplast for synthesis of isoprenoids via the 1-deoxy-D-xylulose 5-phosphate/ 2-*C*-methyl-D-erythritol 4-phosphate pathway. Fosmidomycin acts as a competitive inhibitor of the first committed enzyme in this pathway and its efficacy is decreased when glycolytic flux is increased (44). b) Synchronised WT- and Δ*Pf*PGP-infected RBC were treated with different doses of fosmidomycin for 72 hours and the growth percentage was determined by flow cytometry following SYTO 61 labelling. Data are presented as the mean ± SEM from three independent experiments performed on different days.

To confirm that the metabolic dysregulation of glycolysis and the oxidative pentose phosphate pathway observed in the Δ*Pf*PGP parasite was due to the elevation of 4-PE and not 2-phospho-D-lactate, we incubated WT parasites with D- and L-lactate and performed untargeted LC-MS profiling (Fig. S7). D-lactate incubation led to a selective increase in the phospholactate pool whereas the 4-PE pool remained unchanged. The only other metabolite altered was the D/L-lactate peak itself, indicating that the observed reduction of DHAP and increase in ribose-5-P (Fig. 1c) are most likely the result of 4-PE effects on parasite metabolism.

## Discussion

All eukaryotic and prokaryotic cells express members of the HAD family of metabolite phosphatases, although the function of these proteins *in vivo* are poorly defined. There is increasing evidence that these enzymes have important roles in regulating intracellular levels of a range of phosphorylated intermediates and metabolic fluxes in cells. In this study, we provide evidence that the *P. falciparum* HAD family member, *Pf*PGP, has at least two functions. First, it participates in a novel metabolic cycle involving the phosphorylation and dephosphorylation of D-lactate. Second, it is required to detoxify 4-PE, an allosterically-active metabolic side-product. Targeted deletion of *Pf*PGP resulted in the accumulation of both 2-phospho-D-lactate and 4-PE, attenuated growth of *P. falciparum* asexual blood stages, and increased sensitivity to the antimalarial drug, fosmidomycin. Our results identify important differences in the function of PGP in *P. falciparum* compared to other eukaryotes, consistent with a greater dependence of these parasites on non-transcriptional metabolic regulation.

The PGPs were initially identified in plants as enzymes involved in converting phosphoglycolate, a side product of photorespiration, to glycolate (29). The subsequent conversion of glycolate to 3-phosphoglycerate and its catabolism in the Calvin-Benson cycle increases the efficiency of plant photorespiration by ∼ 25%. More recently, studies on the role of PGP in yeast and animal cells have indicated that these enzymes have broader substrate preferences, acting on metabolites such as glycerol-3-phosphate, 2-phospho-L-lactate and 4-PE (17, 18). Our metabolomic analyses indicate that *Pf*PGP has a similar substrate specificity to the yeast/animal PGPs, although with important differences. In particular, *Pf*PGP appears to act predominantly on the D-enantiomer of 2-phospholactate rather than the L-enantiomer as proposed to occur in animals. L-lactate is the major end-product of glycolysis and 2-phospho-L-lactate is thought to be a minor side product of the glycolytic enzyme, pyruvate kinase, which normally converts pyruvate to L-lactate (17). In contrast, in *P. falciparum* and other eukaryotes that lack a D-lactate-dehydrogenase, D-lactate is produced exclusively via the methylglyoxal pathway. We provide evidence that the 2-phospho-D-lactate detected in wildtype, Δ*Pf*GloI and Δ*Pf*PGP mutants is likely derived from this pathway. Specifically, incubation of wildtype or Δ*Pf*GloI mutant parasites with methylglyoxal led to a concomitant increase in 2-phospho-D-lactate levels. Similarly, incubation of wildtype infected RBC with D-lactate, but not L-lactate, elevates intracellular levels of 2-phospho-D-lactate. The latter result indicates that 2-phospho-D-lactate is synthesized directly from D-lactate, via the action of an as yet unidentified kinase, rather than being a side product of pyruvate kinase.

Interestingly, targeted deletion of the cytoplasmic isoform of GloI, the first enzyme in the methylglyoxal pathway, had little effect on the production of 2-phospho-D-lactate. *P. falciparum* express a second GloI enzyme localized to the apicoplast (30, 31), which could sustain production of D-lactate and 2-phospho-D-lactate in the ΔGloI mutant. Alternatively, or in addition, the RBC host retains a methylglyoxal pathway, which could convert parasite- and host-derived methylglyoxal to D-lactate. The presence of the host pathway may also explain why the parasite glyoxalases are non-essential, although loss of *gloI* results in significant attenuation of intracellular parasite growth over a long period of time (6 replication cycles). Overall, these findings suggest that *P. falciparum* converts D-lactate generated by the parasite (and potentially host) methylglyoxal pathway(s), to 2-phospho-D-lactate via an as yet unidentified kinase. The 2-phospho-D-lactate is subsequently dephosphorylated by *Pf*PGP to regenerate D-lactate. As D-lactate is not secreted by the parasite, *Pf*PGP may contribute to an ATP-consuming futile metabolic cycle that is directly connected to glycolytic flux (Fig. 6).

**Figure 6.**
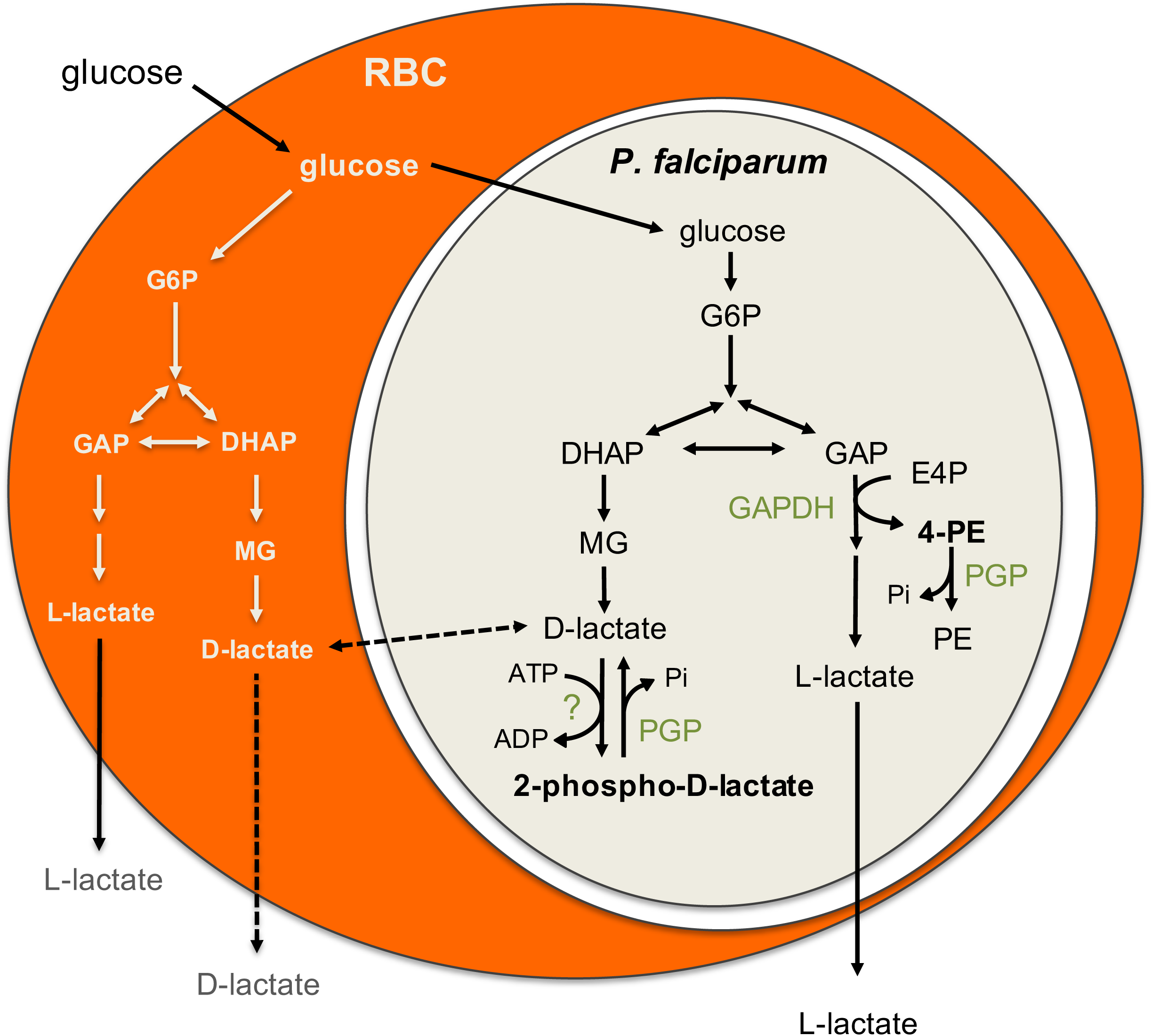
*Pf*PGP is a key metabolic repair enzyme. *P. falciparum* asexual blood stages are heavily dependent on glucose and glycolysis for ATP synthesis, redox balance and production of essential anabolic precursors. PGP is involved in dephosphorylating several metabolites, including 2-phospho-D-lactate and 4-PE, which are produced *in vivo* through the action of unspecified kinase(s) or glycolytic enzymes (GAPDH). These metabolites allosterically regulate glycolytic and PPP enzymes (characteristically involved in catalyzing reactions with ene-diolate intermediates).

What is the potential function of the D-lactate/ 2-phospho-D-lactate cycle in *P. falciparum*? It has recently been shown that 2-phospho-L-lactate (the major phospholactate species in animal cells) inhibits the bi-functional glycolytic enzymes, PFKFB1-4. The enzymes have both phospho-fructo-kinase (PFK) and fructose-2,6-biphosphatase activities and regulate the intracellular levels of the potent PFK allosteric regulator fructose-2,6-bisphosphate (17). Strikingly, *P. falciparum* lacks a PFKFB homologue, and *Pf*PFK is insensitive to conserved metabolic inhibitors/activators (13), highlighting differences in the way these parasites regulate central carbon metabolism. The metabolic futile cycle catalyzed by PGP may have a direct role in maintaining cellular levels of ATP and in preventing excessive glycolytic flux under conditions of nutrient (glucose) excess. While it is difficult to directly estimate the capacity of this cycle to modulate intracellular ATP levels, it is noteworthy that nearly all of the D-lactate produced by infected RBC is retained intracellularly, despite a 100-fold increase in glycolytic flux (as indicated by glucose uptake and L-lactate secretion) (5). This finding suggests that most/all of the D-lactate produced via the parasite methylglyoxal pathway contributes to this cycle. Furthermore, enhanced flux through this pathway, induced by supplementation of infected RBC with D-lactate, led to reduced growth of asexual stages, possibly by increasing ATP cycling. Collectively, these data indicate that the D-lactate/ 2-phospho-D-lactate cycle is very active in these parasite stages and has a significant capacity to modulate ATP levels.

In addition to contributing to the D-lactate/ 2-phospho-D-lactate cycle, *Pf*PGP has a role in regulating intracellular levels of 4-PE. In yeast and animal cells, 4-PE is thought to be generated by the glycolytic enzyme, GAPDH, acting upon the non-standard substrate erythrose-4-P (an intermediate of the pentose phosphate pathway) (17). 4-PE is a potent negative regulator of the enzyme 6-phosphogluconate dehydrogenase in *in vitro* assays (17, 32). In animal PGP- and yeast Pho13- null mutants, the accumulation of 4-PE was proposed to lead to inhibition of the oxidative arm of the PPP (17). In *P. falciparum*, loss of *Pf*PGP resulted in a 9-fold (± 1.6) accumulation of 4-PE indicating that it has a similar role in asexual blood stages. However, in contrast to the situation in yeast and animal cells, significant inhibition of *Pf*6-PGD activity by 4-PE was only observed at 100 μM *in vitro*. In contrast, the *ex vivo* ^13^C-1,2-glucose labelling experiments indicated increased flux through the oxidative PPP in the Δ*Pf*PGP mutant line. These analyses suggest that accumulation of 4-PE is insufficient to inhibit 6-PGD under normal *in vivo* growth conditions. It remains to be determined whether 4-PE inhibits other glycolytic/PPP enzymes in *P. falciparum* that have an ene-diolate intermediate, including triosephosphate isomerase (26, 27) or glucose-6-phosphate isomerase (33). Inhibition of either enzyme could contribute to the reduced glycolytic flux (inferred by the increased sensitivity of Δ*Pf*PGP mutant to fosmidomycin) or cause the increased flux into the oxidative PPP in the Δ*Pf*PGP mutant and explain the growth defect observed for the Δ*Pf*PGP mutant.

This work highlights the important role of HAD enzymes in regulating *P. falciparum* central carbon metabolism. The role of two other *P. falciparum* HAD enzymes have also recently been examined (15, 16). HAD1, the first member of this class to be functionally characterized in *P. falciparum*, promiscuously dephosphorylates a range of glycolytic intermediates *in vitro*, including triose-, pentose- and hexose-phosphates. Loss-of-function mutations in this protein lead to increased parasite resistance to fosmidomycin, and it was proposed that HAD1 has a role in negatively regulating glycolytic flux by removing glycolytic intermediates (15). Similarly, HAD2 dephosphorylates a range of glycolytic intermediates *in vitro*, and is linked to negative regulation of glycolysis (16). A common theme for all three *P. falciparum* HAD proteins characterized to date is their participation in ATP-depleting futile cycles through either depletion of high energy intermediates (HAD1/HAD2) or ATP-dependent cycles of phosphorylation/dephosphorylation (*Pf*PGP). Although these futile cycles appear energetically wasteful, they are comparable to other ATP-dependent cycles, such as protein phosphorylation/dephosphorylation and/or constitutive turnover of mRNA and protein. Metabolic regulation via futile cycling has the potential to be more responsive to subtle changes in carbon sources such as glucose and allows parasites to adapt to changing conditions faster than can be achieved with other forms of metabolic regulation (e.g. transcriptional regulation).

This study highlights the utility of metabolomic approaches in identifying new metabolic pathways and regulatory mechanisms in evolutionarily divergent eukaryotic parasites. A significant proportion of the genes in *Plasmodium* remain uncharacterized and a significant proportion of metabolites detected in comprehensive LC-MS and GC-MS profiling studies have yet to be structurally defined. Our study indicates that unanticipated complexity in cellular metabolism can arise as a result of enzymatic side reactions and/or chemical modification of canonical metabolites, leading to the evolution of new enzyme activities and pathways. We expect that further dissection of the side activities of the major enzymes in intermediary metabolism in *P. falciparum*, and associated repair enzymes will provide new insights into the evolution of novel metabolic regulatory processes in these parasites. Such advances will serve an important role in the identification of new, druggable targets for improved malaria therapeutics.

## Materials and methods

### CRISPR and pTEOE plasmid constructs

GloI and PGP knock-out constructs were cloned using the CRISPR-Cas9 system (23). Briefly, the guide RNA (gRNA) and two homology arms flanking the human dihydrofolate reductase (DHFR) cassette were cloned sequentially into the pL7-CS plasmid. The gRNAs were synthetised and cloned into pL7 using BtgZI restriction sites. Each homology arms sequence was amplified by PCR (CloneAmp - Clontech) from freshly prepared NF54 genomic DNA (Isolate II genomic DNA kit - Bioline) and inserted into pL7 at specific restriction sites (homology arm 1: SpeI/AflII – homology arm 2: EcoRI/NcoI) using InFusion cloning (Clontech). pUF1-Cas9 was used unmodified (23). Plasmid stocks were prepared from maxipreps (Macherey-Nagel). All sequences-of-interest were confirmed by Sanger sequencing (AGRF). The primers used for cloning are presented in Table S2.

For the localization of *Pf*PGP, the coding sequence of *Pf*PGP was amplified by Phusion PCR (NEB) from freshly prepared NF54 genomic DNA and inserted into the pTEOE-GFP plasmid (34) using XhoI and AvrII restriction sites and InFusion cloning (Clontech). To stably integrate the overexpression construct randomly into the genome, the pHTH helper plasmid expressing the *piggyBac* transposase system was used (35). Plasmid stocks were prepared from midipreps (Macherey-Nagel). All sequences-of-interest were confirmed by Sanger sequencing (AGRF). Primers used for cloning are presented in Table S2.

### *P. falciparum* parasite culture and transfections

*P. falciparum* 3D7 and NF54 parasites were cultured in O+ human red blood cells (Australian Red Cross) in RPMI-HEPES–Glutamax (Gibco Life technologies) with 0.25% albumax (Gibco Life technologies), 5% human serum (Australian Red Cross), 10 mM glucose, 50 μM hypoxanthine and gentamycin (Sigma) at 1-4% hematocrit at 37°C in 5% CO_2_, 1% O_2_, Nitrogen balance (Coregas). Cultures were monitored by Giemsa smears. Sorbitol (36) or magnetic enrichment (3) were used to maintain synchronous cultures. Cultures were regularly assessed for mycoplasma contamination (Mycoalert – Lonza) and all experiments reported here are on mycoplasma-free lines.

For knock-out constructs, NF54 parasites were transfected at ring stage using 75 μg of each plasmid (pL7 and pUF-1-Cas9). For overexpression constructs, NF54 parasites were transfected at ring stage using 100 μg of pTEOE and 50 μg of pHTH as previously described (37). DNA/Cytomix were electroporated into red blood cells at 310V, 950 uF as previously described (38). Parasites were maintained on 5 nM WR99210 (Sigma-Aldrich) selection at all times. Knock-out parasites were also selected on 1.5 μM DSM-1 (MR4) for the first five days post-transfection. 38 μM 5-fluorocytosine (negative selection) was added to recovered transfectants and healthy rings returned after 2-6 days (39).

Validation of successful integration was confirmed by PCR on knock-out-NF54 genomic DNA (Bioline), using primer pairs that were specific to the integrated cassette and the genomic DNA. Primers used for this purpose are presented in Table S2.

### *P. falciparum* parasite growth assay

The parasitemia of synchronised ring cultures (0.8% haematocrit, 0.8% parasitemia) was assessed by flow cytometry each day for 13 days and identical dilutions between wildtype and knock-out infected RBC were applied regularly. Nucleic acids were stained with SYTO 61 (Invitrogen) and parasitemia was measured on a FACS Canto^TM^ II (BD Biosciences). Data were analysed using the FlowJo software (BD Biosciences). Raw counts were normalized to the day 13 WT values and the dilutions made throughout the experiment were corrected for.

### D-lactate *in vitro* assay

The concentration of D-lactate excreted from uninfected- and NF54 infected- RBC was measured using a D-lactate assay kit (Cayman chemical). All conditions were incubated for 26-29 hours (rings to trophozoite stage) in complete RPMI media. Media was collected, deproteinised and measured following the manufacturer’s instructions. After incubation for 30 minutes at 37°C, the plate was read at 590 nm (Ex. 544 nm) on a FluoStar Omega plate reader (BMG Labtech) and analysed with the Omega data software.

### *in vitro* assays

Infected-RBC were magnetically enriched at trophozoite stage (MAGNEX Cell Separator (Colebrook Bioscience); >95% parasitemia) and lysed in 0.1% saponin buffer. After three PBS washes and counting of isolated parasites, cell pellets were snap frozen in liquid nitrogen and stored at -80°C. Pellets were resuspended in 200 μL per 1.10^8^ cells of pH7.4 lysis buffer (5 mM HEPES, 2 mM DTT, protease inhibitor). Enzymatic reactions were setup in a 50:50 ratio cell lysate:reaction buffer. For measuring glyoxalase activity, the reaction buffer was composed of 100 mM Tris HCl, 5 mM NH_4_Cl, 2 mM MgCl_2_, 2 mM ATP, 2 mM methylglyoxal (MG) and 2 mM reduced glutathione. The control reaction consisted of the reaction buffer without MG (- MG condition) and used as the baseline for data normalization. Enzymatic reactions were setup at 37°C and samples were collected in technical duplicates at 0, 5, 10, 30 and 60 minutes.

6-PGD activity was measured as described above using a reaction buffer composed of 100 mM Tris HCl, 5 mM NH_4_Cl, 2 mM MgCl_2_, 1 mM 6-phospho gluconate, 1 mM NADP and 1mM 4-phosphoerythronate (as annotated). Enzymatic reactions were setup at 37°C, incubated for 60 minutes and collected in technical duplicates at 0 and 60 minutes. Samples were extracted for GC-MS analysis (see details below).

### GC-MS sample extraction and derivatisation

All *in vitro* GC-MS samples were extracted on ice with 100 μL chloroform and 400 μL 3:1 methanol:H_2_O containing 1 nM scyllo-inositol (internal standard). At the end of the sample collection, 200 μL H_2_O was added to each tube, samples were mixed thoroughly and centrifuged at 14 000 x g for 5 minutes at room temperature. The aqueous phase was transferred into a new tube, ready for GC-MS drying.

Samples were dried in a SpeedVac (Savant). Pellets were washed in 600 μL 90% methanol. Thirty μL were transferred into mass spectrometry tubes (1/20 dilution) and dried (SpeedVac). 50μL of pure methanol were used for an additional wash to remove any trace water and samples were dried in a SpeedVac. Samples were derivatised with 20 μL of 20 mg/mL methoxyamine (Sigma Aldrich) made up in pyridine (Sigma Aldrich) and left at room temperature overnight. The next morning, 20 μL of a ready-to-use N,O-bis(trimethylsilyl) trifluoroacetamide (BSTFA) + 1 % trimethylchlorosilane (TMCS) solution (Supelco) was added to each sample. Metabolites were separated as described previously (40) using a BD5 capillary column (J&W Scientific, 30 m x 250 μM x 0.25 μM) on a Hewlett Packard 6890 system (5973 EI-quadrupole MS detector). Briefly, the oven temperature gradient was 70 °C (1 minute); 70 °C to 295 °C at 12.5 °C/minute, 295 °C to 320 °C at 25 °C/minute; 320 °C for 2 minutes. MS data was acquired using scan mode with a *m/z* range of 50-550, threshold 150 and scan rate of 2.91 scans/second. GC retention time and mass spectra were compared with authentic standards analysed in the same batch for metabolite identification.

### ^13^C glucose labelling and LC-MS analysis

Synchronised trophozoite cultures were magnetically enriched using a MAGNEX Cell Separator (Colebrook Bioscience). Purified cells (5×10^7^ cells/sample) were incubated for 30 minutes at 37°C in either RPMI containing 11 mM ^13^C_1,2_-glucose (Cambridge Isotopes) (= fully labelled) or 11 mM ^13^C-U-glucose mixed 1:1 with complete RPMI containing 11 mM unlabelled glucose (Sigma). Samples were centrifuged at 14 000 x g for 30 seconds, washed in 1 mL ice-cold PBS and pellets were extracted for GC-MS analysis for ^13^C_1,2_-glucose labelled samples (as described above) or extracted for LC-MS analysis (^13^C-U-glucose samples).

LC-MS samples were resuspended in 100 μL of 80% acetonitrile (Burdick & Jackson) containing the internal standard 1 μM 4-^13^C,^15^N-aspartate. After centrifugation at 14 000 x g for 5 minutes at 4°C, supernatants were transferred into mass spectrometry vials. LC-MS analysis was performed as described previously (40), with the following modifications. Metabolite samples were separated on a SeQuant ZIC-pHILIC column (5 μM, 150 x 4.6 mm, Millipore) using a binary gradient with a 1200 series HPLC system (Agilent), with solvent A being water with 20 mM ammonium carbonate and solvent B 100% acetonitrile. The gradient ran linearly (at 0.3 mL/minute) from 80-20% solvent B from 0.5 to 15 minutes, then 20-5% between 15 and 20 minutes, before returning to 80% at 20.5 minutes and kept at 80% solvent B until 29.5 minutes. MS detection was performed on an Agilent Q-TOF mass spectrometer 6545 operating in negative ESI mode. The scan range was 80-1200 *m/z* between 2 and 25 minutes at 0.9 spectra/second. An internal reference ion solution continually run (isocratic pump at 0.2 mL/minute) throughout the chromatographic separation to maintain mass accuracy. Other LC parameters were: autosampler temperature 4 °C, injection volume 10 μL and data were collected in centroid mode with Mass Hunter Workstation software (Agilent).

The untargeted profiling of Δ*Pf*PGP-infected RBC compared to WT-infected RBC was performed as described above and the effect of L- and D-lactate on parasite metabolism was assessed by incubating purified infected RBC in complete RPMI containing +/- 10 mM L- and D-lactate (Sigma) for one hour at 37°C and analysed via LC-MS as described above.

### Mass spectrometry data analysis

GC-MS data was processed using ChemStation (Agilent) or the in-house software package DExSI (41) and metabolites of interest were compared to authentic standards. LC-MS.d files were converted to .mzXML files using MS convert and analysed using MAVEN (42). Following alignment, peaks were extracted with a mass tolerance of <10 ppm. Untargeted comparative profiling was performed to generate a list of *m/z* features of interest. The *m/z* value of each peak of interest was then queried against the METLIN metabolite database (METLIN reference) for only M-H adducts with a 10 ppm mass tolerance. Peaks of interest were positively identified using their exact mass and retention time (compared to a standards library of 150 compounds ran the same day).

### Synthesis of sodium phospholactate

Pearlman’s catalyst (20 % Pd(OH)_2_/C, 135 mg) was added to a solution of phospho(enol)pyruvic acid monosodium salt monohydrate (Sigma, 40 mg, 192 μmol) in THF/MeOH/AcOH 1:1:1 (3 mL) and the mixture was stirred at room temperature under an atmosphere of H_2_ (40 bar) for 48 hours. The reaction mixture was diluted with MeOH (20 mL) then filtered through Celite, and evaporated at 10 mbar on a rotary evaporator at 40 °C. The residue was dissolved in deionized water, and the solution was passed through a C18 Sep-Pak. The eluent was evaporated at 10 mbar on a rotary evaporator at 40 °C, and the residue was dried at 10^-3^ mbar at room temperature for 24 hours, affording D/L-phospholactate monosodium salt hydrate (40 mg) as a viscous oil. HRMS (ESI-TOF) calculated (calcd) for C_3_H_4_O_6_P^-^ [M – Na]^-^, 168.9907 *m*/*z*, found 168.9917.

### Synthesis of cyclohexylammonium 4-phospho-D-erythronate

Br_2_ saturated H_2_O (616 μL, ∼35 mg/mL, 135 μmol) was added to a stirred mixture of D-erythrose-4-phosphate (Sigma, 12.0 mg, 54.0 μmol) in aqueous Na_2_CO_3_ (400 μL, 0.405 M, 162 μmol) at room temperature. After one hour, excess bromine was removed by sparging for 2 hours with N_2_. The resultant mixture was passed down a column containing Amberlite Ag50 (acid form, 2 mL bed volume, ∼1.7 mM H^+^ per mL), which was rinsed with H_2_O (3 × 1.5 mL). Cyclohexylamine (39 μL, 0.324 mmol) was added to the eluted product, and the volatiles were removed by rotary evaporation and further drying under an N_2_ stream overnight. The crude material was purified on a Shimadzu 2020 LC-MS instrument, using an ACE Excel 5 Super C18 column (150 mm x 2.1 mm) with 0.1 % formic acid (solvent A) and methanol (solvent B). A linear gradient was performed from 80 % to 10 % across 15 minutes and elution of 4-PE was determined using the theoretical exact mass of 4-PE. Separation of 4-PE from contaminants was monitored using the MS in full scan mode and UV detection. Verification of the purified 4-PE was performed by re-running the collected aliquot on the instrument and confirming no other masses or UV peaks were observed (above background).

### Fluorescence microscopy

Glass coverslips were incubated with 0.1 mg/mL PHAE (Sigma-Aldrich) for 30 minutes at 37°C in a humid chamber. After 3 washes in PBS, infected-RBC (3% haematocrit) were incubated for 10 minutes on the coverslip. Nuclei were stained using 2 μg/mL DAPI for 10 minutes. Slides were mounted in *p*-phenylenediamine antifade. Images were taken on a Delta Vision Elite restorative widefield deconvolution Imaging system (GE Healthcare) using a 100x UPLS Apo objective (1.4 NA, Olympus) lens under oil immersion. The following Emission/ Excitation filter sets were used: DAPI Ex 390/18, Em 435/48 – FITC Ex 475/28, Em 523/26. Images were deconvoluted using Softworx 5.0 (GE Healthcare) and analysed using ImageJ (NIH).

### Protein analysis by western blotting

Trophozoite parasites were isolated by addition of 0.05% saponin in the culture media and spun down at 3 750 x g for 5 minutes. Pellets were washed twice in ice-cold PBS containing complete protease inhibitors (Roche) and resuspended in PBS or RIPA buffers containing protease inhibitors. RIPA-buffer samples were incubated on ice for 10 min and centrifuged at 16 000 x g for 10 minutes. The supernatant was collected and placed into a fresh tube. The saponin pellet (PBS) and the RIPA (supernatant) samples were mixed with Bolt 4X LDS and 10X reducing agent (Invitrogen) and heated at 85°C for 10 minutes prior SDS-PAGE on 4-12% BisTris gels separated in 1X MOPS running buffer (Invitrogen). Proteins were transferred onto nitrocellulose membranes using the iBlot 2 transfer system (Invitrogen) and membranes were blocked in 3.5% skim milk for at least one hour at room temperature. Membranes were probed with mouse anti-GFP (1:1000 - Roche) and rabbit anti-GAPDH (1:1000 - (43)) primary antibodies. Secondary antibodies were horseradish-peroxidase conjugated: goat anti-mouse and anti-rabbit (1:20 000 - Promega). Membranes were incubated with Clarity ECL substrate (Bio-Rad) and imaged on a ChemiDoc MP system (BioRad).

## Acknowledgments

The authors thank Dr Marion Hliscs for technical assistance with preliminary work, Dr Natalie Spillman, Dr Stuart Ralph and Prof Geoff McFadden for use of reagents and helpful discussions. We thank Metabolomics Australia, Bio21 Institute, for the use of GC- and LC-MS equipment. We thank the Australian Red Cross service for blood donations.

## Supporting information

**Figure S1. *Pf*PGP shares homology with other described PGP enzymes** CLUSTAL alignment of *P. falciparum* PGP (PF3D7_0715000) with known sequences of yeast Pho13 (YDL236W), mouse AUM (Q8CHP8) and human PGP (NC_000016). The four characteristic HAD motifs (21) framed in red (I to IV) appear to be highly conserved between the compared species.

**Figure S2. *Pf*PGP is a cytosolic protein and molecular characterisation of Δ*Pf*PGP parasites** a) Western blot of PGP-GFP- (PGP) and WT-infected RBC lysates probed with anti-GFP and anti-GAPDH (loading control). Samples were extracted in either PBS or RIPA buffer. This blot is representative of three independent experiments. b) Schematic of the Δ*Pf*PGP cloning strategy. Legends: CDS = coding sequence; UTR = untranslated regions; hDHFR = resistance cassette; (k)bp = (kilo) base pairs. Features: striped boxes = homology arms; short black line = guide RNA; dotted line between arrows = fragment used for genetic confirmation of the knock-out. c) Genetic confirmation of the disruption of the *pgp* gene by PCR on genomic DNA. A 1 kb DNA ladder was used as reference.

**Figure S3. Identification of 2-phospho-D-lactate in wildtype parasites** a) 2-Phospholactate GC-MS spectrum with fingerprint signature. b) 3D7 WT-infected RBC were incubated with D-lactate (1 mM) or left untreated (iRBC condition) and increasing concentrations of a pure 2-phospholactate standard was spiked into cell extracts. The y-axis represents the arbitrary ion counts and the x-axis represents the retention time in minutes for the phospholactate peak.

**Figure S4. Molecular characterisation of Δ*Pf*GloI parasites** a) Schematic of the Δ*Pf*GloI cloning strategy. Legends: CDS = coding sequence; UTR = untranslated regions; hDHFR = resistance cassette; (k)bp = (kilo) base pairs. Features: striped boxes = homology arms; short black line = guide RNA; dotted line between arrows = fragment used for genetic confirmation of the knock-out. b) Genetic confirmation of the disruption of the *gloI* gene by PCR on genomic DNA. The single band at 1.1 kb in the knockout line only matches the expected 1 145 bp PCR fragment. A 1 kb DNA ladder was used as reference.

**Figure S5. Levels of 4-phosphoerythronate are not affected by addition of exogenous D/L lactate** 4-PE levels were measured by GC-MS in Δ*Pf*PGP- vs WT- infected RBC upon 1-hour incubation at 37°C with 2 mM D- or L-lactate. Data are presented as fold change to the WT - no lactate condition. The mean values ± SEM from three to four independent replicates performed on different days are presented.

**Figure S6. The inhibition of 6-phosphogluconate dehydrogenase is specific to 4-phosphoerythronate** 6-Phosphogluconate dehydrogenase *in vitro* activity was tested on saponin-isolated trophozoite parasites by measurement of the ribulose-5-P product by GC-MS. Parasite lysates were incubated with (left to right): no 4-PE, 1 mM phospholactate (+Plac), 1 mM erythronate (+Ery), no NADP and no reaction buffer (lysate). Results are normalised to the no 4-PE condition (100%). Data are presented as the mean ± SEM from three independent experiments performed on different days.

**Figure S7. The increase in 2-phospho-D-lactate is not responsible for metabolic dysregulations observed in Δ*Pf*PGP parasite cultures** WT-infected RBC were subjected to a 1-hour treatment with 10 mM L- or D-lactate and metabolically compared to a no-treatment condition (control treated) by untargeted LC-MS. Scatter plots of *m/z* ion counts are presented. Following both D- and L-lactate exposure, the intracellular lactate pool was significantly elevated (*P* < 0.05 Benjamini corrected). Phospholactate was significantly elevated only following D-lactate incubation (*P* < 0.05 Benjamini corrected). Each scatter plot represents averaged data from three (L-lactate) and four (D-lactate) independent biological replicates.

**Table S1**. **Repertoire of the metabolites-of-interest and analytical GC-MS features** Highlight of the quantified ion and retention time used for identifying specific metabolites in all GC-MS-based assays presented in this manuscript.

**Table S2. Primer list** Primers are presented by section. For the CRISPR and pTEOE cloning sections, InFusion primers were designed. Upper cases are nucleotides that are part of the plasmid backbone, lower cases are nucleotides that are part of the gene-of-interest, underlined nucleotides correspond to restriction sites, and for GloI_HA1_rev_AflII the nucleotides in blue are mutated nucleotides.

